# Cross-laboratory analysis of brain cell type transcriptomes with applications to interpretation of bulk tissue data

**DOI:** 10.1101/089219

**Authors:** B. Ogan Mancarci, Lilah Toker, Shreejoy J Tripathy, Brenna Li, Brad Rocco, Etienne Sibille, Paul Pavlidis

## Abstract

Establishing the molecular diversity of cell types is crucial for the study of the nervous system. We compiled a cross-laboratory database of mouse brain cell type-specific transcriptomes from 36 major cell types from across the mammalian brain using rigorously curated published data from pooled cell type microarray and single cell RNA-sequencing studies. We used these data to identify cell type-specific marker genes, discovering a substantial number of novel markers, many of which we validated using computational and experimental approaches. We further demonstrate that summarized expression of marker gene sets in bulk tissue data can be used to estimate the relative cell type abundance across samples. To facilitate use of this expanding resource, we provide a user-friendly web interface at Neuroexpresso.org.

**Significance Statement:** Cell type markers are powerful tools in the study of the nervous system that help reveal properties of cell types and acquire additional information from large scale expression experiments. Despite their usefulness in the field, known marker genes for brain cell types are few in number. We present NeuroExpresso, a database of brain cell type specific gene expression profiles, and demonstrate the use of marker genes for acquiring cell type specific information from whole tissue expression. The database will prove itself as a useful resource for researchers aiming to reveal novel properties of the cell types and aid both laboratory and computational scientists to unravel the cell type specific components of brain disorders.

## Introduction

Brain cells can be classified based on features such as their primary type (e.g. neurons vs. glia), location (e.g. cortex, hippocampus, cerebellum), electrophysiological properties (e.g. fast spiking vs. regular spiking), morphology (e.g. pyramidal cells, granule cells) or the neurotransmitter/neuromodulator they release (e.g. dopaminergic cells, serotonergic cells, GABAergic cells). Marker genes, genes that are expressed in a specific subset of cells, are often used in combination with other cellular features to define different types of cells (Hu et al., 2014; Margolis et al., 2006) and facilitate their characterization by tagging the cells of interest for further studies (Handley et al., 2015; Lobo et al., 2006; Tomomura et al., 2001). Marker genes have also found use in the analysis of whole tissue “bulk” gene expression profiling data, which can be challenging to interpret due to the difficulty to determine the source of the observed expressional change. For example, a decrease in a transcript level can indicate a regulatory event affecting the expression level of the gene, a decrease in the number of cells expressing the gene, or both. To address this issue, computational methods have been proposed to estimate cell type specific proportion changes based on expression patterns of known marker genes (Chikina et al., 2015; Newman et al., 2015; Westra et al., 2015; Xu et al., 2013). Finally, marker genes are obvious candidates for having cell type specific functional roles.

An ideal cell type marker has a strongly enriched expression in a single cell type in the brain. However, this criterion can rarely be met, and for many purposes, cell type markers can be defined within the context of a certain brain region; namely, a useful marker may be specific for the cell type in one region but not necessarily in another region or brain-wide. For example, the calcium binding protein parvalbumin is a useful marker of both fast spiking interneurons in the cortex and Purkinje cells in the cerebellum (Celio and Heizmann, 1981; Kawaguchi et al., 1987). Whether the markers are defined brain-wide or in a region-specific context, the confidence in their specificity is established by testing their expression in as many different cell types as possible. This is important because a marker identified by comparing just two cell types might turn out to be expressed in a third, untested cell type, reducing its utility.

During the last decade, targeted purification of cell types of interest followed by gene expression profiling has been applied to many cell types in the brain. Such studies, targeted towards well-characterized cell types, have greatly promoted our understanding of the functional and molecular diversity of these cells (Cahoy et al., 2008; Chung et al., 2005; Doyle et al., 2008). However, individual studies of this kind are limited in their ability to discover specific markers as they often analyse only a small subset of cell types (Shrestha et al., 2015; Okaty et al., 2009; Sugino et al., 2006) or have limited resolution as they group subtypes of cells together (Cahoy et al., 2008). Recently, advances in technology have enabled the use of single cell transcriptomics as a powerful tool to dissect neuronal diversity and derive novel molecular classifications of cells (Poulin et al., 2016). However, with single cell analysis the classification of cells to different types is generally done post-hoc, based on the clustering similarity in their gene expression patterns. These molecularly defined cell types are often uncharacterized otherwise (e.g. electrophysiologically, morphologically), challenging their identification outside of the original study and understanding their role in normal and pathological brain function. A notable exception is the single cell RNA-seq study of Tasic et al. (2016) analysing single labelled cells from transgenic mouse lines to facilitate matching of the molecularly defined cell types they discover to previously identified cell types. We hypothesized that aggregating cell type specific studies that analyse expression profiles of cell types previously defined in literature, a more comprehensive data set more suitable for marker genes could be derived.

Here we report the analysis of an aggregated cross-laboratory dataset of cell type specific expression profiling experiments from mouse brain, composed both of pooled cell microarray data and single cell RNA-seq data. We used these data to identify sets of brain cell marker genes more comprehensive than any previously reported, and validated the markers genes in external mouse and human single cell datasets. We further show that the identified markers are applicable for the analysis of human brain and demonstrate the usage of marker genes in the analysis of bulk tissue data via the summarization of their expression into “marker gene profiles” (MGPs), which can be cautiously interpreted as correlates of cell type proportion. Finally, we made both the cell type expression profiles and marker sets available to the research community at neuroexpresso.org.

## Materials and methods

Figure 1A depicts the workflow and the major steps of this study. All the analyses were performed in R version 3.3.2; the R code and data files can be accessed through neuroexpresso.org (RRID: SRC_015724) or directly from <u>https://github.com/oganm/neuroexpressoAnalysis.</u>

**Figure 1:**
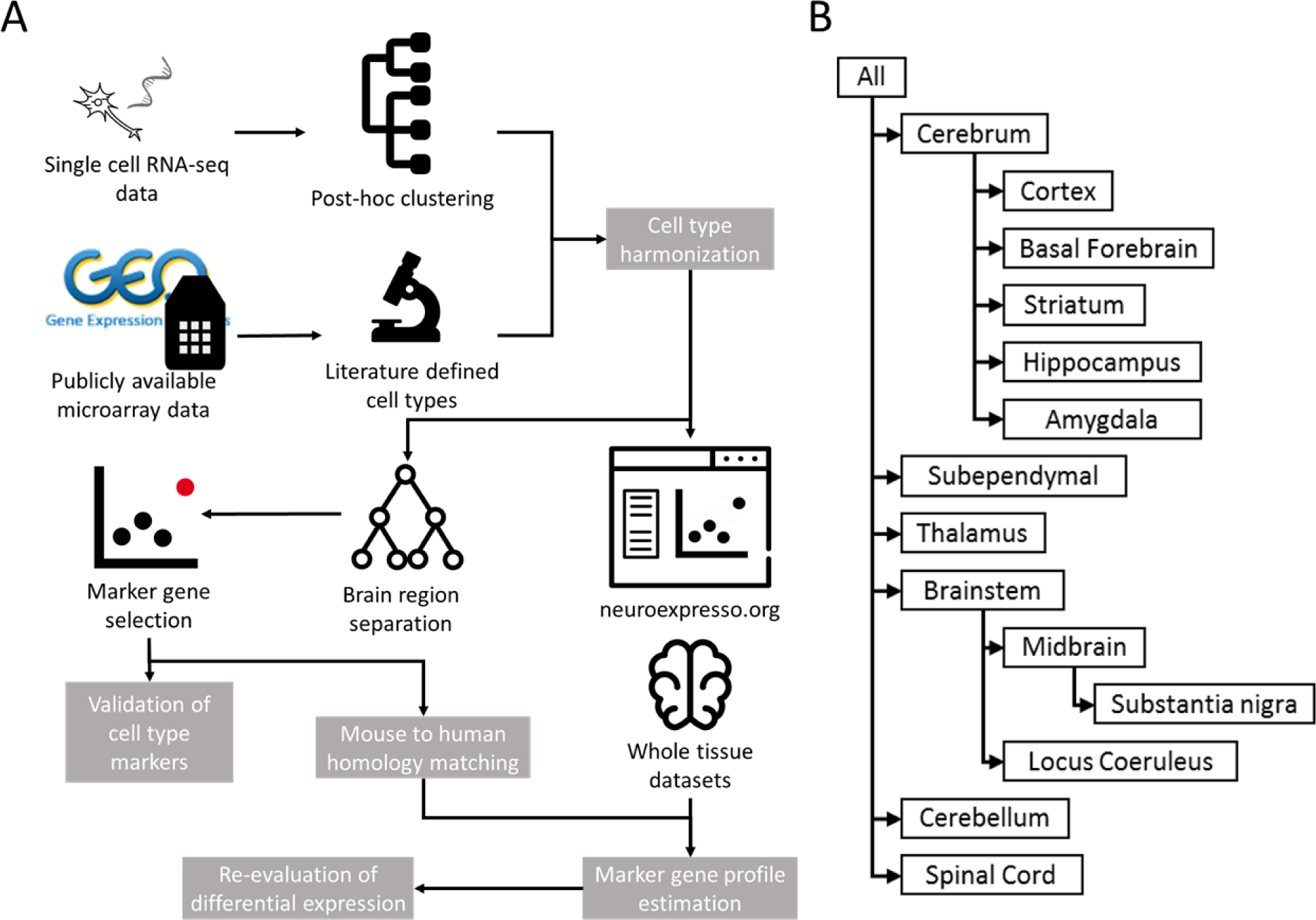
Mouse brain cell type specific expression database compiled from publicly available datasets. **(A)** Workflow of the study. Cell type specific expression profiles are collected from publicly available datasets and personal communications. Acquired samples are grouped based on cell type and brain region. Marker genes are selected per brain region for all cell types. Marker genes are biologically and computationally validated and used in estimation of cell type proportions. **(B)** Brain region hierarchy used in the study. Samples included in a brain region based on the region they were extracted from. For instance, dopaminergic cells isolated from the midbrain were included when selecting marker genes in the context of brainstem and whole brain. Microglia extracted from whole brain isolates were added to all brain regions.

### Pooled cell type specific microarray data sets

We began with a collection of seven studies of isolated cell types from the brain, compiled by Okaty et al. (2011). We expanded this by querying PubMed (<u>http://www.ncbi.nlm.nih.gov/pubmed)</u> and Gene Expression Omnibus (GEO) (<u>http://www.ncbi.nlm.nih.gov/geo/)</u> (RRID: SCR_007303) (Barrett et al., 2013; Edgar et al., 2002) for cell type-specific expression datasets from the mouse brain that used Mouse Expression 430A Array (GPL339) or Mouse Genome 430 2.0 Array (GPL1261) platforms. These platforms were our focus as together, they are the most popular platforms for analysis of mouse samples and are relatively comprehensive in gene coverage, and using a reduced range of platforms reduced technical issues in combining studies. Query terms included names of specific cell types (e.g. astrocytes, pyramidal cells) along with blanket terms such as “brain cell expression” and “purified brain cells”. Only samples derived from postnatal (> 14 days), wild type, untreated animals were included. Datasets obtained from cell cultures or cell lines were excluded due to the reported expression differences between cultured cells and primary cells (Cahoy et al., 2008; Halliwell, 2003; Januszyk et al., 2015). We also considered RNA-seq data from pooled cells (2016; Zhang et al., 2014) but because such data sets are not available for many cell types, including it in the merged resource was not technically feasible without introducing biases (though we were able to incorporate a single-cell RNA-seq data set, described in the next section). While we plan to incorporate more pooled cell RNA-seq data in the future, for this study we limited their use to validation of marker selection. As a first step in the quality control of the data, we manually validated that each sample expressed the gene that was used as a marker for purification of the corresponding cell type in the original publication (expression greater than median expression among all gene signals in the dataset), along with other well established marker genes for the relevant cell type (e.g. Pcp2 for Purkinje cells, Gad1 for GABAergic interneurons). We next excluded contaminated samples, namely, samples expressing established marker genes of non-related cell types in levels comparable to the cell type marker itself (for example neuronal samples expressing high levels of glial marker genes), which lead to the removal of 21 samples. In total, we have 30 major cell types compiled from 24 studies represented by microarray data (summarized in Table 1); a complete list of all samples including those removed is available from the authors).

**Table 1:**
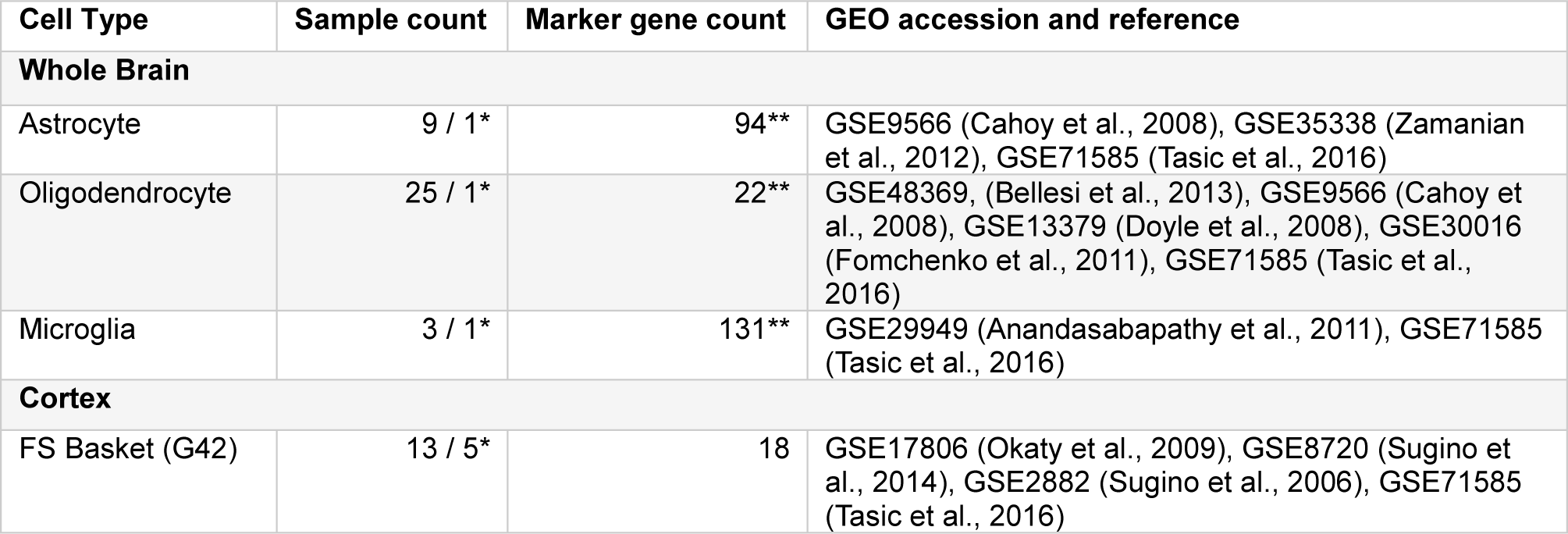
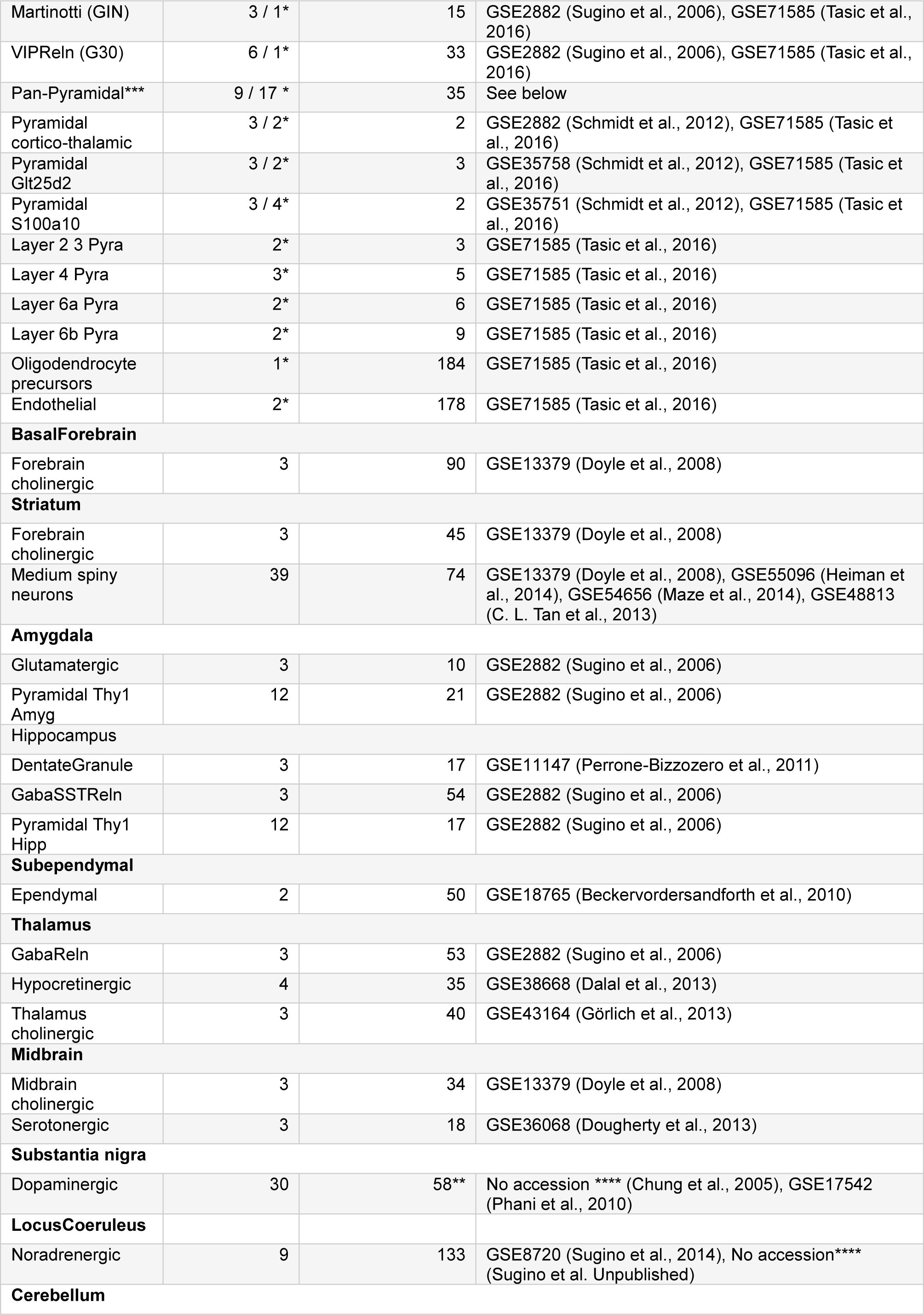
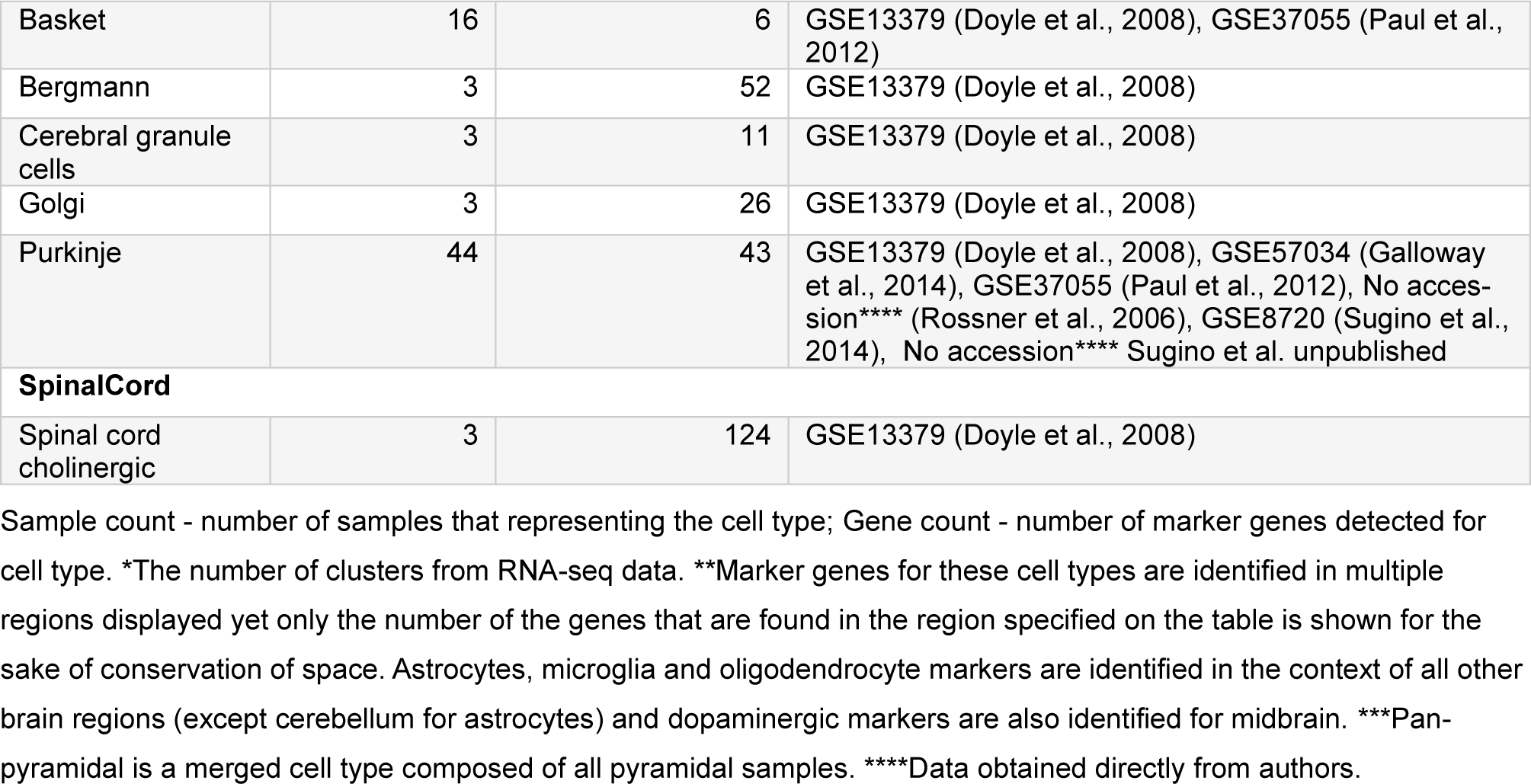
Cell types in NeuroExpresso database.

### Single cell RNA-seq data

The study of cortical single cells by Tasic et al. (2016) includes a supplementary file (Supplementary Table 7 in Tasic et al. (2016)) linking a portion of the molecularly defined cell clusters to known cell types previously described in the literature. Using this file, we matched the cell clusters from Tasic et al. with pooled cortical cell types represented by microarray data (<u>Table 2</u>). For most cell types represented by microarray (e.g. glial cells, Martinotti cells), the matching was based on the correspondence information provided by Tasic et al. (2016). However, for some of the cell clusters from Tasic et al. (2016), the cell types were matched manually, based on the description of the cell type in the original publication (e.g., cortical layer, high expression of a specific gene). For example, Glt25d2^+^ pyramidal cells from Schmidt et al. (2012), described by the authors as “layer 5b pyramidal cells with high Glt25d2 and Fam84b expression” were matched with two cell clusters from Tasic et al. - “L5b Tph2” and “L5b Cdh13”, 2 of the 3 clusters described as Layer 5b glutamatergic cells by Tasic et al., since both of these clusters represented pyramidal cells from cortical layer 5b and exhibited high level of the indicated genes. Cell clusters identified in Tasic et al. that did not match to any of the pooled cell types were integrated into to the combined data if they fulfilled the following criteria: 1) They represented well-characterized cell types and 2) we could determine with high confidence that they did not correspond to more than one cell type represented by microarray data. <u>Table 2</u> contains information regarding the matching between pooled cell types from microarray data and cell clusters from single cell RNA-seq data from Tasic et al.

**Table 2:** Matching single cell RNA sequencing data from Tasic to well defined cell types.

| Microarray cell type | Tasic et al. cell cluster | Matching method | NeuroExpresso cell type name |
| --- | --- | --- | --- |
| Astrocyte | Astro Gja1 | Direct match | Astrocyte |
| Microglia | Micro Ctss | Direct match | Microglia |
| Oligodendrocyte | Oligo Opalin | Direct match | Oligodendrocyte |
| FS Basket (G42) | Pvalb Gpx3, Pvalb Rspo2, Pvalb Wt1, Pvalb Obox3, Pvalb Cpne5 | Definition: fast spiking pval positive interneurons | FS Basket (G42) |
| Martinotti (GIN) | Sst Cbln4 | Direct match | Martinotti (GIN) |
| VIPReIn (G30) | Vip Sncg | Unique Vip and Sncg expression, high Sncg expression in microarray cell type | VIPReIn (G30) |
| Pyramidal Glt25d2 | L5b Tph2, L5b Cdh13 | Definition: Glt25d2 positive Fam84b positive | Pyramidal Glt25d2 |
| Pyramidal S100a10 | L5a Hsd11b1, L5a Batf3, L5a Tcerg1l, L5a Pde1c | Definition: S100a10 expressing cells from layer 5a | Pyramidal S100a10 |
| Pyramidal CrtThalamic | L6a Car12, L6a Syt17 | Direct match | Pyramidal Crt-Thalamic |
| --- | Endo Myl9, Endo Tbc1d4 | New cell type | Endothelial |
| --- | OPC Pdgfra | New cell type | Oligodendrocyte precursors |
| --- | L4 Ctxn3, L4 Scnn1a, L4 Arf5 | New cell type | Layer 4 Pyra |
| --- | L2 Ngb, L2/3 Ptgs2 | New cell type | Layer 2 3 Pyra |
| --- | L6a Mgp, L6a Sla | New cell type | Layer 6a Pyra |
| --- | L6b Serpinb11, L6b Rgs12 | New cell type | Layer 6b Pyra |

List of molecular cell types identified by Tasic et al. and their corresponding cell types in NeuroExpresso. Matching method column defines how the matching was performed. Direct matches are one to one matching between the definition provided by Tasic et al. for the molecular cell types and definition provided by microarray samples. For “Definition” matches, description of the cell type in the original source is used to find molecular cell types that fit the definition. VIPReIn – Vip Sncg matching was done based on unique Sncg expression in VIPReIn cells in the microarray data. New cell types are well defined cell types that have no counterpart in microarray data.

In total, the combined database contains expression profiles for 36 major cell types, 10 of which are represented by both pooled cell microarray and single cell RNA-seq data, and five which are represented by single cell RNA-seq only (summarized in <u>Table 2</u>). Due to the substantial differences between microarray and RNA-seq technologies, we analysed these data separately (see next sections). For visualization only, in neuroexpesso.org we rescaled the RNA-seq data to allow them to be plotted on the same axes. Details are provided on the web site.

### Grouping and re-assignment of cell type samples

When possible, samples were assigned to specific cell types based on the descriptions provided in their associated publications. When expression profiles of closely related cell types were too similar to each other and we could not find sufficient number of differentiating marker genes meeting our criteria, they were grouped together into a single cell type. For example, A10 and A9 dopaminergic cells had no distinguishing markers (provided the other cell types presented in the midbrain region) and were grouped as “dopaminergic neurons”. In the case of pyramidal cells, while we were able to detect marker genes for pyramidal cell subtypes, they were often few in numbers and most of them were not represented on the human microarray chip (Affymetrix Human Exon 1.0 ST Array) used in the downstream analysis. As a result, calculation of marker gene profiles in human bulk tissue would not feasible for majority of these cell types. To combat this, we created two gene lists, one created by considering pyramidal subtypes as separate cell types, and another where pyramidal subtypes are pooled into a pan-pyramidal cell type. Due to the scarcity of markers for pyramidal subtypes, we only consider the pan-pyramidal cell type in our downstream analysis. However, we still kept the pyramidal subtypes separate during marker gene selection (described below) for the non-pyramidal cell types to help ensure marker specificity.

Since our focus was identifying markers specific to cell types within a given brain region, samples were grouped based on the brain region from which they were isolated, guided by the anatomical hierarchy of brain regions (Figure 1B). Brain sub-regions (e.g. locus coeruleus) were added to the hierarchy if there were multiple cell types represented in the sub-region. An exception to the region assignment process are glial samples. Since these samples were only available from either cortex or cerebellum regions or extracted from whole brain, the following assignments were made: Cerebral cortex-derived astrocyte and oligodendrocyte samples were included in the analysis of other cerebral regions as well as thalamus, brainstem and spinal cord. Bergmann glia and cerebellum-derived oligodendrocytes were used in the analysis of cerebellum. The only microglia samples available were isolated from whole brain homogenates and were included in the analysis of all brain regions.

### Selection of cell type markers

Marker gene sets (MGSs) were selected for each cell type in each brain region, based on fold change and clustering quality (see below). For cell types that are represented by both microarray and single cell data (cortical cells), two sets of MGSs were created and later merged as described below. Since there is no generally accepted definition of “marker gene”, our goal was to identify markers that were sufficiently specific and highly expressed to be useful in computational settings, but also likely to be of interest for potential laboratory applications. Thus, our threshold selections were guided in part by the expression patterns of previously well-established markers as well as our intended applications.

Marker genes were selected for each brain region based on the following steps:

1. For RNA-seq data, each of the relevant clusters identified in Tasic et al. was considered as a single sample, where the expression of each gene was calculated by taking the mean RPKM values of the individual cells representing the cluster. <u>Table 2</u> shows which clusters represent which cell types.
2. Expression level of a gene in a cell type was calculated by taking the mean expression of all replicate samples originating from the same study and averaging the resulting values across different studies per cell type.
3. The quality of clustering was determined by “mean silhouette coefficient” and “minimal silhouette coefficient” values (where silhouette coefficient is a measure of group dissimilarity ranged between - 1 and 1 (Rousseeuw, 1987)). Mean silhouette coefficient was calculated by assigning the samples representing the cell type of interest to one cluster and samples from the remaining cell types to another, and then calculating the mean silhouette coefficient of all samples. The minimal silhouette coefficient is the minimal value of mean silhouette coefficient when it is calculated for samples representing the cell type of interest in comparison to samples from each of the remaining cell types separately. The two measures where used to ensure that the marker gene robustly differentiates the cell type of interest from other cell types. Silhouette coefficients were calculated with the “silhouette” function from the “cluster” R package version 1.15.3 (Maechler et al., 2016), using the expression difference of the gene between samples as the distance metric.
4. A background expression value was defined as expression below which the signal cannot be discerned from noise. Different background values are selected for microarray (6 – all values are log_2_ transformed) and RNA-seq (0.1) due to the differences in their distribution.

Based on these metrics, the following criteria were used:

1. A threshold expression level was selected to help ensure that the gene’s transcripts will be detectable in bulk tissue. Genes with median expression level below this threshold were excluded from further analyses. For microarrays, this threshold was chosen to be 8. Theoretically, if a gene has an expression level of 8 in a cell type, and the gene is specific to the cell type, an expression level of 6 would be observed if 1/8^th^ of a bulk tissue is composed of the cell type. As many of the cell types in the database are likely to be as rare as or rarer than 1/8^th^, and 6 is generally close to background for these data, we picked 8 as a lower level of marker gene expression. For RNA-seq data, we selected a threshold of 2.5 RPKM, which in terms of quantiles corresponds to the microarray level of 8.
2. If the expression level in the cell type of interest is higher than 10 times the background threshold, there must be at least a 10-fold difference from the median expression level of the remaining cell types in the region. If the expression level in the cell type is less than 10 times the background, the expression level must be higher than the expression level of every other cell type in that region. This criterion was added because below this expression level, for a 10-fold expression change to occur, the expression median of other cell types needs be lower than the background. Values below the background signal that do not convey meaningful information but can prevent potentially useful marker genes from being selected.
3. The mean silhouette coefficient for the gene must be higher than 0.5 and minimum silhouette coefficient must the greater than zero for the associated cell type.
4. The conditions above must be satisfied only by a single cell type in the region.

To ensure robustness against outlier samples, we used the following randomization procedure, repeated 500 times: One third (rounded) of all samples were removed. For microarray data, to prevent large studies from dominating the silhouette coefficient, when studies representing the same cell types did not have an equal number of samples, N samples were picked randomly from each of the studies, where N is the smallest number of samples coming from a single study. A gene was selected if it qualified our criteria in more than 95% of all permutations.

Our next step was combining the MGSs created from the two expression data types. For cell types and genes represented by both microarray and RNA-seq data, we first looked at the intersection between the MGSs. For most of the cell types, the overlap between the two MGSs was about 50%. We reasoned that this could be partially due to numerous “near misses” in both data sources. Namely, since our method for marker gene selection relies on multiple steps with hard thresholds, it is very likely that some genes were not selected simply because they were just below one of the required thresholds. We thus adopted a soft intersection: A gene was considered as a marker if it fulfilled the marker gene criteria in one data source (pooled cell microarray or single cell RNA-seq), and its expression in the corresponding cell type from the other data source was higher than in any other cell type in that region. For example, Ank1 was originally selected as a marker of FS Basket cells based on microarray data, but did not fulfil our selection criteria based on RNA-seq data. However, the expression level of Ank1 in the RNA-seq data is higher in FS Basket cells than in any other cell type from this data source, and thus, based on the soft intersection criterion, Ank1 is considered as a marker of FS Basket cells in our final marker gene set. For genes and cell types that were only represent by one data source, the selection was based on this data source only.

It can be noted that some previously described markers (such as Prox1 for dentate granule cells) are absent from our marker gene lists. In some cases, this is due to the absence the genes from the microarray platforms used, while in other cases the genes failed to meet our stringent selection criteria. Final marker gene lists are available at <u>http://www.chibi.ubc.ca/supplement-to-mancarci-et-al-neuroexpresso/.</u> Human homologues of mouse genes were defined by NCBI HomoloGene (<u>ftp://ftp.ncbi.nih.gov/pub/HomoloGene/build68/homologene.data</u>).

### Microglia enriched genes

Microglia expression profiles differ significantly between activated and inactivated states and to our knowledge, the samples in our database represent only the inactive state (Holtman et al., 2015). In order to acquire marker genes with stable expression levels regardless of microglia activation state, we removed the genes differentially expressed in activated microglia based on Holtman et al. (2015). This step resulted in removal of 408 out of the original 720 microglial genes in cortex (microarray and RNA-seq lists combined) and 253 of the 493 genes in the context of other brain regions (without genes from single cell data). Microglial marker genes which were differentially expressed in activated microglia are referred to as Microglia_activation and Microglia_deactivation (up or down-regulated, respectively) in the marker gene lists provided.

### S100a10^+^ pyramidal cell enriched genes

The paper (Schmidt et al., 2012) describing the cortical S100a10^**+**^ pyramidal cells emphasizes the existence of non-neuronal cells expressing S100a10^+^. Schmidt et al. therefore limited their analysis to 7,853 genes specifically expressed in neurons and advised third-party users of the data to do so as well. Since a contamination caveat was only concerning microarray samples from Schmidt et al. (the only source of S100a10^+^ pyramidal cells in microarray data), we removed marker genes selected for S100a10^+^ pyramidal cells based on the microarray data if they were not among the 7,853 genes indicated in Schmidt et al. We also removed S100a10 itself since based on the author’s description it was not specific to this cell type. In total, 36 of the 47 S100a10 pyramidal genes originally selected based on microarray data were removed in this step. Of note, none of the removed genes were selected as a marker of S100a10 cell based on RNA-seq data.

### Dentate granule cell enriched genes

We used data from (Cembrowski et al., 2016) (Hipposeq – RRID: SCR_015730) for validation and refinement of dentate granule markers (as noted above these data are not currently included in Neuroexpresso for technical reasons). FPKM values were downloaded (GEO accession GSE74985) and log_2_ transformed. Based on these values, dentate granule marker genes were removed if their expression in Hipposeq data (mean of dorsal and ventral granule cells) was lower than other cell types represented in this dataset. In total, 15 of the 39 originally selected genes that were removed in this step.

### In situ hybridization

Male C57BL/6J (RRID: IMSR_JAX:0000664) mice aged 13-15 weeks at time of sacrifice were used (n=5). Mice were euthanized by cervical dislocation and then the brain was quickly removed, frozen on dry ice, and stored at - 80°C until sectioned via cryostat. Brain sections containing the sensorimotor cortex were cut along the rostral-caudal axis using a block advance of 14 μm, immediately mounted on glass slides and dried at room temperature (RT) for 10 minutes, and then stored at - 80°C until processed using multi-label fluorescent in situ hybridization procedures.

Fluorescent in situ hybridization probes were designed by Advanced Cell Diagnostics, Inc. (Hayward, CA, USA) to detect mRNA encoding Cox6a2, Slc32a1, and Pvalb. Two sections per animal were processed using the RNAscope^®^ 2.5 Assay as previously described (Wang et al., 2012). Briefly, tissue sections were incubated in a protease treatment for 30 minutes at RT and then the probes were hybridized to their target mRNAs for 2 hours at 40°C. The sections were exposed to a series of incubations at 40°C that amplifies the target probes, and then counterstained with NeuroTrace blue-fluorescent Nissl stain (1:50; Molecular Probes) for 20 minutes at RT. Cox6a2, Pvalb, and Slc32a1 were detected with Alexa Fluor^®^ 488, Atto 550 and Atto 647, respectively.

Data were collected on an Olympus IX83 inverted microscope equipped with a Hamamatsu Orca-Flash4.0 V2 digital CMOS camera using a 60× 1.40 NA SC oil immersion objective. The equipment was controlled by cellSens (Olympus). 3D image stacks (2D images successively captured at intervals separated by 0.25 μm in the z-dimension) that are 1434 × 1434 pixels (155.35 μm × 155.35 μm) were acquired over the entire thickness of the tissue section. The stacks were collected using optimal exposure settings (i.e., those that yielded the greatest dynamic range with no saturated pixels), with differences in exposures normalized before analyses.

Laminar boundaries of the sensorimotor cortex were determined by cytoarchitectonic criteria using NeuroTrace labeling. Fifteen image stacks across the gray matter area spanning from layer 2 to 6 were systematic randomly sampled using a sampling grid of 220 × 220 μm^2^, which yielded a total of 30 image stacks per animal. Every NeuroTrace labeled neuron within a 700 × 700 pixels counting frame was included for analyses; the counting frame was placed in the center of each image to ensure that the entire NeuroTrace labeled neuron was in the field of view. The percentage (± standard deviation) of NeuroTrace labeled cells containing Cox6a2 mRNA (Cox6a2+) and that did not contain Slc32a1 mRNA (Slc32a1-), that contained Slc32a1 but not Pvalb mRNA (Slc32a1+/Pvalb-), and that contained both Slc32a1 and Pvalb mRNAs (Slc32a1+/Pvalb+) were manually assessed.

### Allen Brain Atlas in situ hybridization (ISH) data

We downloaded in situ hybridization (ISH) images using the Allen Brain Atlas API (<u>http://help.brain-map.org/display/mousebrain/API).</u> Assessment of expression patterns was done by visual inspection. If a probe used in an ISH experiment did not show expression in the region, an alternative probe targeting the same gene was sought. If none of the probes showed expression in the region, the gene was considered to be not expressed.

## Validation of marker genes using external single cell data

Mouse cortex single cell RNA sequencing (RNA-seq) data were acquired from Zeisel et al. (2015) (available from <u>http://linnarssonlab.org/cortex/,</u> GEO accession: GSE60361,1691 cells) Human single cell RNA sequencing data were acquired from Darmanis et al. (2015) (GEO accession: GSE67835, 466 cells). For both datasets, pre-processed expression data were encoded in a binary matrix with 1 representing any nonzero value. For all marker gene sets, Spearman’s ρ was used to quantify internal correlation. A null distribution was estimated by calculating the internal correlation of 1000 randomly-selected prevalence-matched gene groups. Gene prevalence was defined as the total number of cells with a non-zero expression value for the gene. Prevalence matching was done by choosing a random gene with a prevalence of +/-2.5% of the prevalence of the marker gene. P-values were calculated by comparing the internal correlation of marker gene set to the internal correlations of random gene groups using Wilcoxon rank-sum test.

## Pre-processing of microarray data

For comparison of marker gene profiles in white matter and frontal cortex, we acquired expression data from pathologically healthy brain samples from Trabzuni et al. (2013) (GEO accession: GSE60862). For estimation of dopaminergic marker gene profiles in Parkinson’s disease patients and controls, we acquired substantia nigra expression data from Lesnick et al. (2007) (GSE7621), Moran et al.(2006) (GSE8397) and Zhang et al.(2005) (GSE20295) studies. Expression data for the Stanley Medical Research Institute (SMRI), which included post-mortem prefrontal cortex samples from bipolar disorder, major depression and schizophrenia patients along with healthy donors, were acquired through <u>https://www.stanleygenomics.org/,</u> study identifier 2.

All microarray data used in the study were pre-processed and normalized with the “rma” function of the “oligo” (RRID: SCR_015729) (Affymetrix gene arrays) or “affy” (RRID: SCR_012835) (Affymetrix 3’IVT arrays) (Carvalho and Irizarry, 2010) R packages. Probeset to gene annotations were obtained from Gemma (Zoubarev et al., 2012) (<u>http://gemma.chibi.ubc.ca/</u>). Probesets with maximal expression level lower than the median among all probeset signals were removed. Of the remaining probesets, whenever several probesets were mapped to the same gene, the one with the highest variance among the samples was selected for further analysis.

Outliers and mislabelled samples were removed when applicable, if they were identified as an outlier in provided metadata, if expression of sex-specific genes did not match the sex provided in metadata (Toker et al., 2016), or if they clustered with data from another tissue type in the same dataset based on genes found to be most differentially expressed between the tissue types. This resulted in the removal of 18/194 samples from Trabzuni et al. (2013), 3/44 samples from expression data from Stanley Medical Research Institute and 3/93 samples from Zhang et al. (2005) dataset.

Samples from pooled cell types that make up the NeuroExpresso database were processed by an in-house modified version of the “rma” function that enabled collective processing of data from Mouse Expression 430A Array (GPL339) and Mouse Genome 430 2.0 Array (GPL1261) which share 22690 of their probesets. As part of the rma function, the samples are quantile normalized at the probe level. However, possibly due to differences in the purification steps used by different studies (Okaty et al., 2011), we still observed biases in signal distribution among samples originating from different studies. Thus, to increase the comparability across studies, we performed a second quantile normalization of the samples at a probeset level before selection of probes with the highest variance. After all processing the final data set included 11564 genes.

### Estimation of marker gene profiles (MGPs)

For each cell type, relevant to the brain region analysed, we used the first principal component of the corresponding marker gene set expression as a surrogate for cell type proportions. This method of marker gene profile estimation is similar to the methodology of multiple previous works that aim to estimate relative abundance of cell types in a whole tissue sample (Chikina et al., 2015; Westra et al., 2015; Xu et al., 2013). Principal component analysis was performed using the “prcomp” function from the “stats” R package, using the “scale = TRUE” option. It is plausible that some marker genes will be transcriptionally differentially regulated under some conditions (e.g. disease state), reducing the correspondence between their expression level with the relative cell proportion. A gene that is thus regulated is expected to have reduced correlation to the other marker genes with expression levels primarily dictated by cell type proportions, which will reduce their loading in the first principal component. To reduce the impact of regulated genes on the estimation process, we removed marker genes from a given analysis if their loadings had the opposite sign to the majority of markers when calculated based on all samples in the dataset and recalculate loadings and components using the remaining genes. This was repeated until all remaining genes had loadings with the same signs. Since the sign of the loadings of the rotation matrix (as produced by prcomp function) is arbitrary, to ease interpretation between the scores and the direction of summarized change in the expression of the relevant genes, we multiplied the scores by - 1 whenever the sign of the loadings was negative. For visualization purposes, the scores were normalized to the range 0-1. Two sided Wilcoxon rank-sum test (“wilcox.test” function from the “stats” package in R, default options) was used to compare between the different experimental conditions.

For estimations of cell type MGPs in samples from frontal cortex and white matter from the Trabzuni study (Trabzuni et al., 2013), results were subjected to multiple testing correction by the Benjamini & Hochberg method (Benjamini and Hochberg, 1995). For the Parkinson’s disease datasets from Moran et al. (2006) and Lesnick et al. (2007), we estimated MGPs for dopaminergic neuron markers in control and PD subjects. Moran et al. data included samples from two sub-regions of substantia nigra. Since some of the subjects were sampled in only one of the sub-regions while others in both, the two sub-regions were analysed separately.

For the SMRI collection of psychiatric patients we estimated oligodendrocytes MGPs based on expression data available through the SMRI website (as indicated above) and compared our results to experimental cell counts from the same cohort of subjects previously reported by Uranova et al. (2004). Figure 7B representing the oligodendrocyte cell counts in each disease group was adapted from Uranova et al. (2004). The data presented in the figure was extracted from Figure 1A in Uranova et al. (2004) using WebPlotDigitizer (<u>http://arohatgi.info/WebPlotDigitizer/app/</u>).

**Figure 7:**
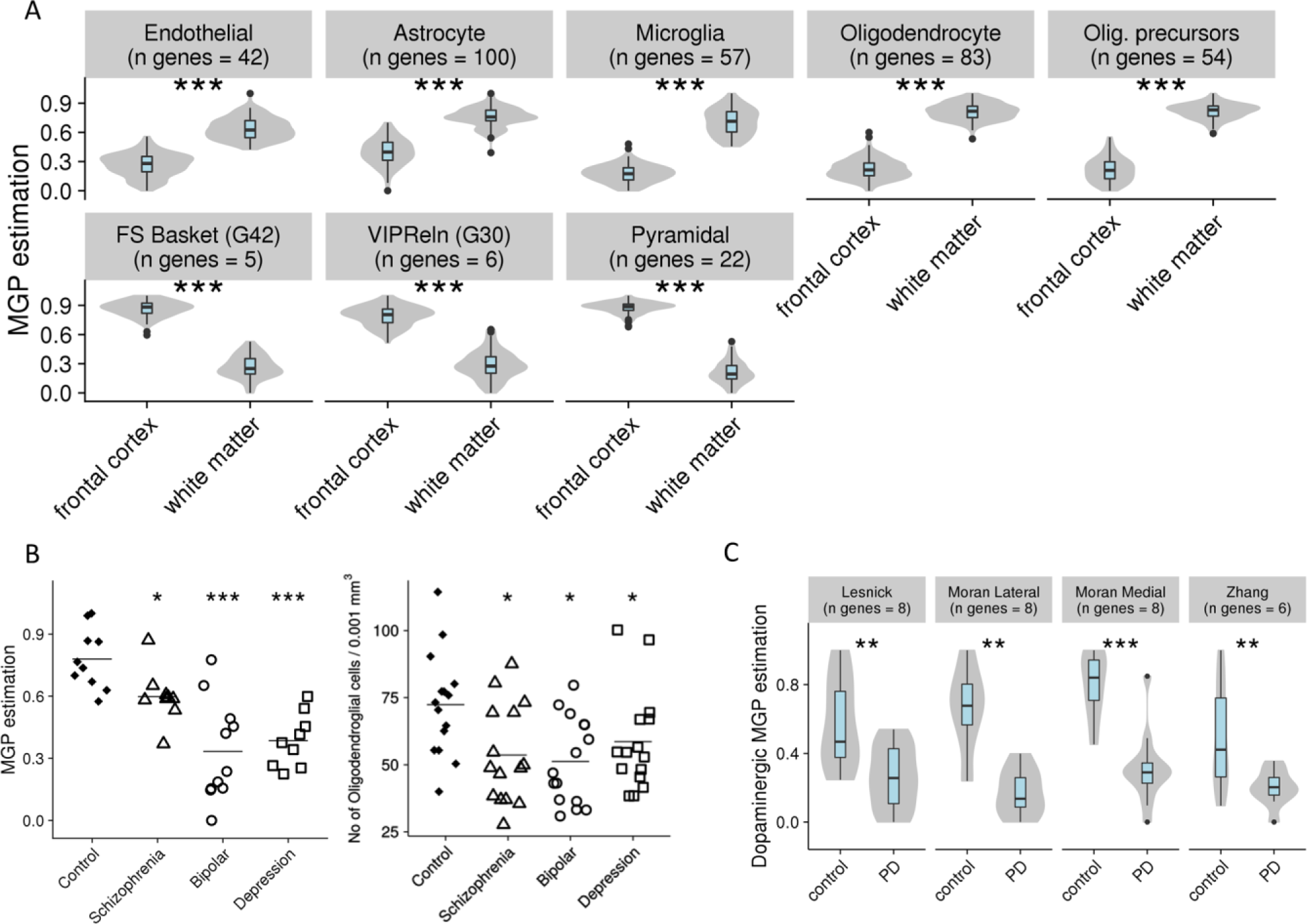
Marker gene profiles reveal cell type specific changes in whole tissue data. **(A)** Estimation of cell type profiles for cortical cells in frontal cortex and white matter. Values are normalized to be between 0 and 1. (***p<0.001). **(B)** Left: Oligodendrocyte MGPs in Stanley C cohort. Right: Morphology based oligodendrocyte counts of Stanley C cohort. Figure adapted from Uranova et al. (2004). **(C)** Estimations of dopaminergic cell MGPs in substantia nigra of controls and Parkinson’s disease patients. Values are relative and are normalized to be between 0 and 1 and are not reflective of absolute proportions (**p <0.01, ***p<0.001).

### Code Accessibility

All code is available as extended data. They are also maintained in the GitHub repositories lister below. Marker gene selection and marker gene profile estimation was performed with custom R functions provided within “markerGeneProfile” R package available on GitHub (<u>https://github.com/oganm/markerGeneProfile</u>). Human homologues of mouse genes were identified using “homologene” R package available on GitHub (<u>https://github.com/oganm/homologene</u>).

Code for data processing and analysis can be found at “neuroExpressoAnalysis” repository available on GitHub (<u>https://github.com/oganm/neuroExpressoAnalysis</u>).

Source code of the neuroexpresso.org we app can be found at “neuroexpresso” repository available on GitHub (<u>https://github.com/oganm/neuroexpresso</u>)

## Results

### Compilation of a brain cell type expression database

A key input to our search for marker genes is expression data from purified pooled brain cell types and single cells. Expanding on work from Okaty et al. (2011), we assembled and curated a database of cell type-specific expression profiles from published data (see Methods, Figure 1A). The database represents 36 major cell types from 12 brain regions (Figure 1B) from a total of 263 samples and 30 single cell clusters. Frontal cortex is represented by both microarray and RNA-seq data, with 5 of the 15 cortical cell types represented exclusively by RNA-seq data. We used rigorous quality control steps to identify contaminated samples and outliers (see Methods). In the microarray dataset, all cell types except for ependymal cells are represented by at least 3 replicates and in the entire database, 14/36 cell types are represented by multiple independent studies (Table 1). The database is in constant growth as more cell type data becomes available. To facilitate access to the data and allow basic analysis we provide a simple search and visualization interface on the web, <u>www.neuroexpresso.org</u> (Figure 2). The app provides means of visualising gene expression in different brain regions based on the cell type, study or methodology, as well as differential expression analysis between groups of selected samples.

**Figure 2:**
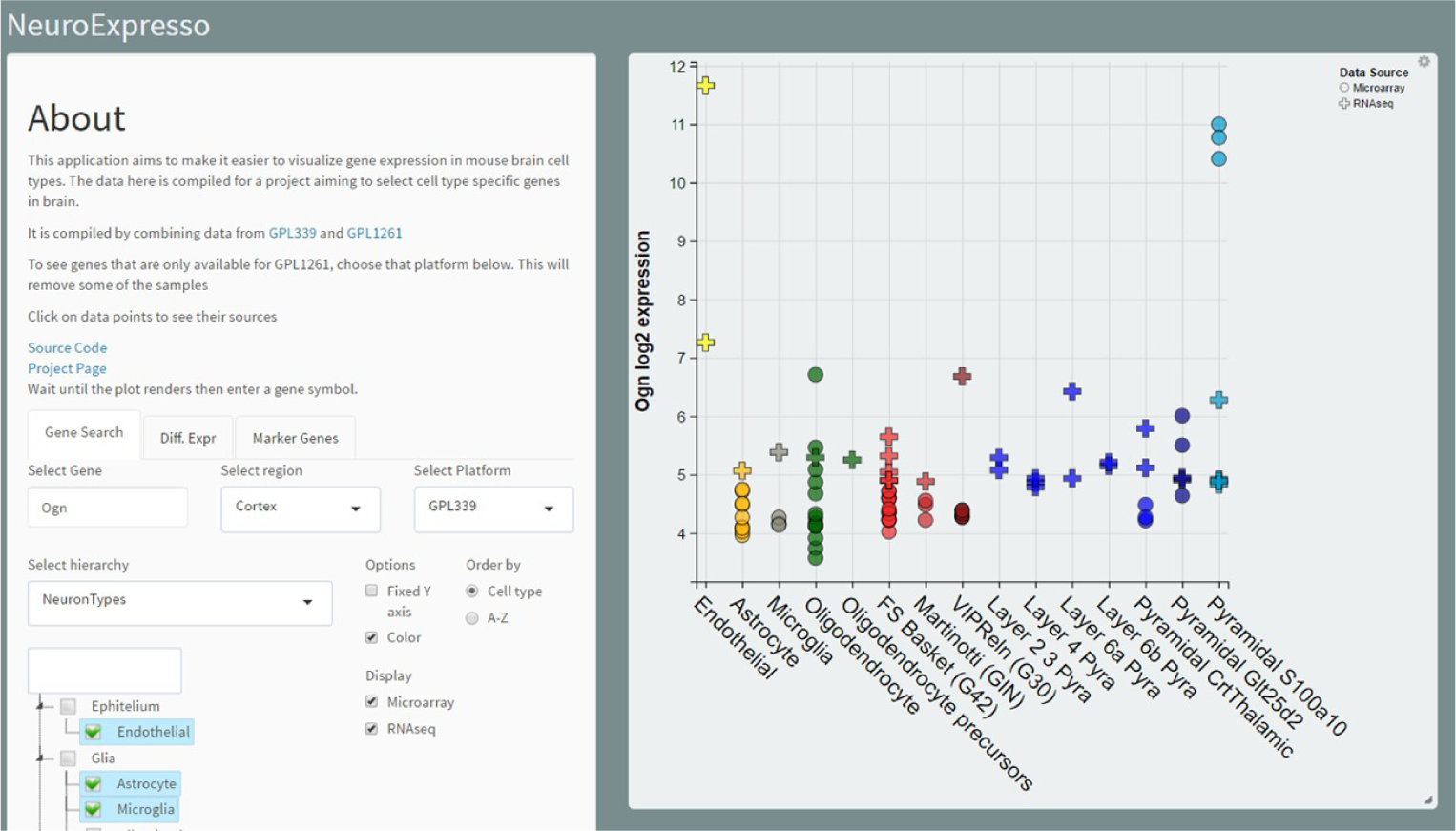
The NeuroExpresso.org web application. The application allows easy visualization of gene expression across cell types in brain regions. Depicted is the expression of cell types from frontal cortex region. Alternatively, cell types can be grouped based on their primary neurotransmitter or the purification type. The application can be reached at <u>www.neuroexpresso.org.</u>

### Identification of cell type enriched marker gene sets

We used the NeuroExpresso data to identify marker gene sets (MGSs) for each of the 36 cell types. An individual MGS is composed of genes highly enriched in a cell type in the context of a brain region (Figure 3A). Marker genes were selected based on a) fold of change relative to other cell types in the brain region and b) a lack of overlap of expression levels in other cell types (see Methods for details). This approach captured previously known marker genes (e.g. Th for dopaminergic cells (Pickel et al., 1976), Tmem119 for microglia (Bennett et al., 2016) (of note, Tmem119 was classified as downregulated in activated microglia in our analysis, corroborating previous reports of Satoh et al. (2016) and Erny et al. (2015)). We also identified numerous new candidate markers such as Cox6a2 for fast spiking parvalbumin (PV)^+^ interneurons. Some marker genes previously reported by individual studies whose data were included in our database, were not selected by our analysis. For example, Fam114a1 (9130005N14Rik), identified as a marker of fast spiking basket cells by Sugino et al. (2006), is highly expressed in oligodendrocytes and oligodendrocyte precursor cells (Figure 3B). These cell types were not considered in the Sugino et al. (2006) study, and thus the lack of specificity of Fam114a1 could not be observed by the authors. In total, we identified 2671 marker genes (3- 186 markers per cell type, (Table 1)). The next sections focus on verification and validation of our proposed markers, using multiple methodologies.

**Figure 3:**
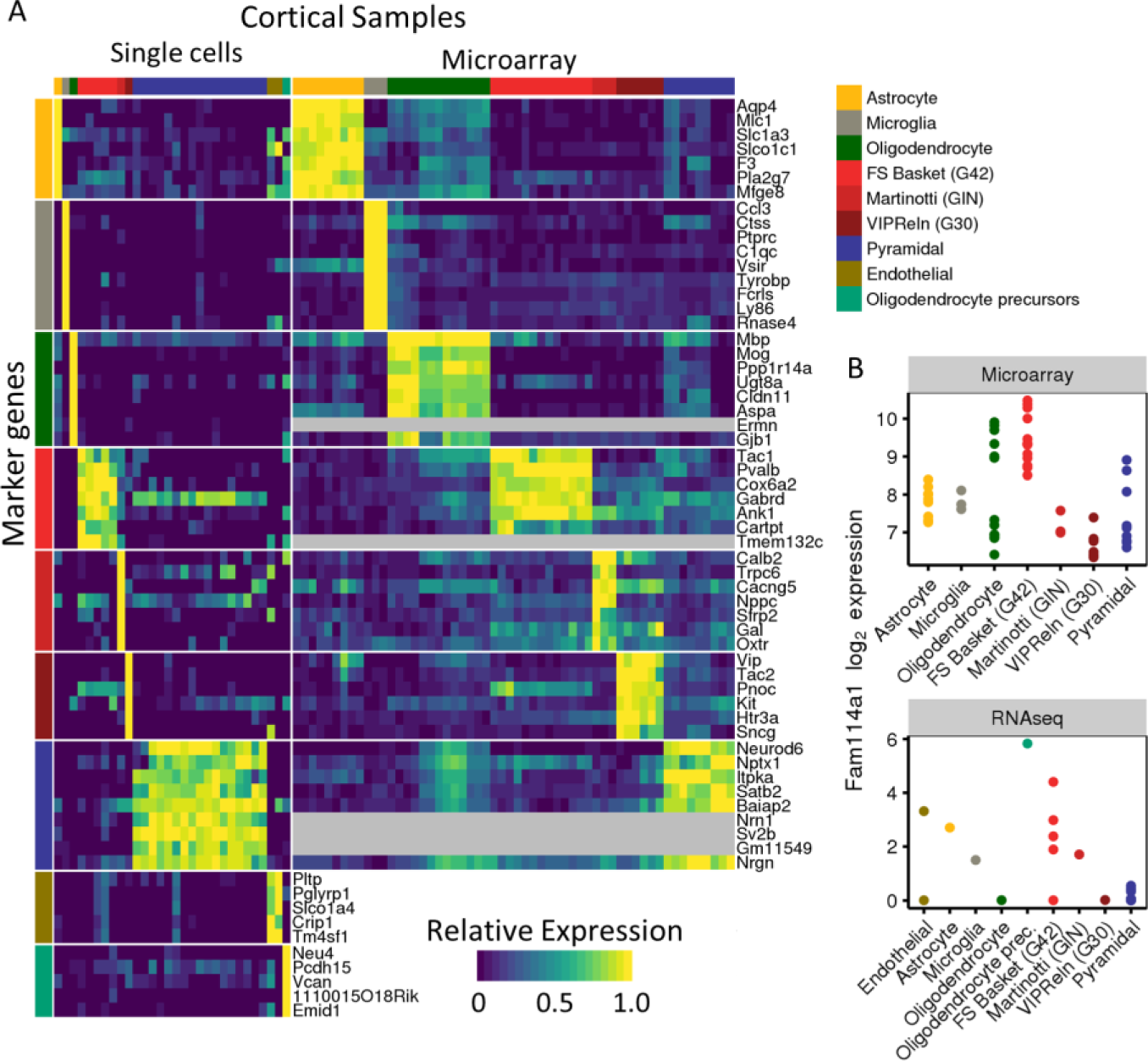
Marker genes are selected for mouse brain cell types and used to estimate cell type profiles. **(A)** Expression of top marker genes selected for cell cortical cell types in cell types represented by RNA-seq (left) and microarray (right) data in NeuroExpresso. Expression levels were normalized per gene to be between 0-1 for each dataset. **(B)** Expression of Fam114a1 in frontal cortex in microarray (left) and RNA-seq (right) datasets. Fam114a1 is a proposed fast spiking basket cell marker. It was not selected as a marker in this study due to its high expression in oligodendrocytes and S100a10 expressing pyramidal cells that were both absent from the original study.

### Verification of markers by in situ hybridization

Two cell types in our database (Purkinje cells of the cerebellum and hippocampal dentate gyrus granule cells) are organized in well-defined anatomical structures that can be readily identified in tissue sections. We exploited this fact to use in situ hybridization (ISH) data from the Allen Brain Atlas (ABA) (<u>http://mouse.brain-map.org)</u> (Sunkin et al., 2013) to verify co-localization of known and novel markers for these two cell types. There was a high degree of co-localization of the markers to the corresponding brain structures, and by implication, cell types (Figure 4A-B). For dentate granule (DG) cell markers, all 16 genes were represented in ABA. Of these, 14 specifically co-localized with known markers (that is, had the predicted expression pattern confirming our marker selection), one marker exhibited non-specific expression and one marker showed no signal. For Purkinje cell markers, 41/43 genes were represented in ABA. Of these, 37 specifically co-localized with known markers, one marker exhibited non-specific expression and three markers showed no signal in the relevant brain structure (Figure 4B). Figure 4A shows representative examples for the two cell types (details of our ABA analysis, including images for all the genes examined and validation status of the genes, are provided in extended data).

**Figure 4:**
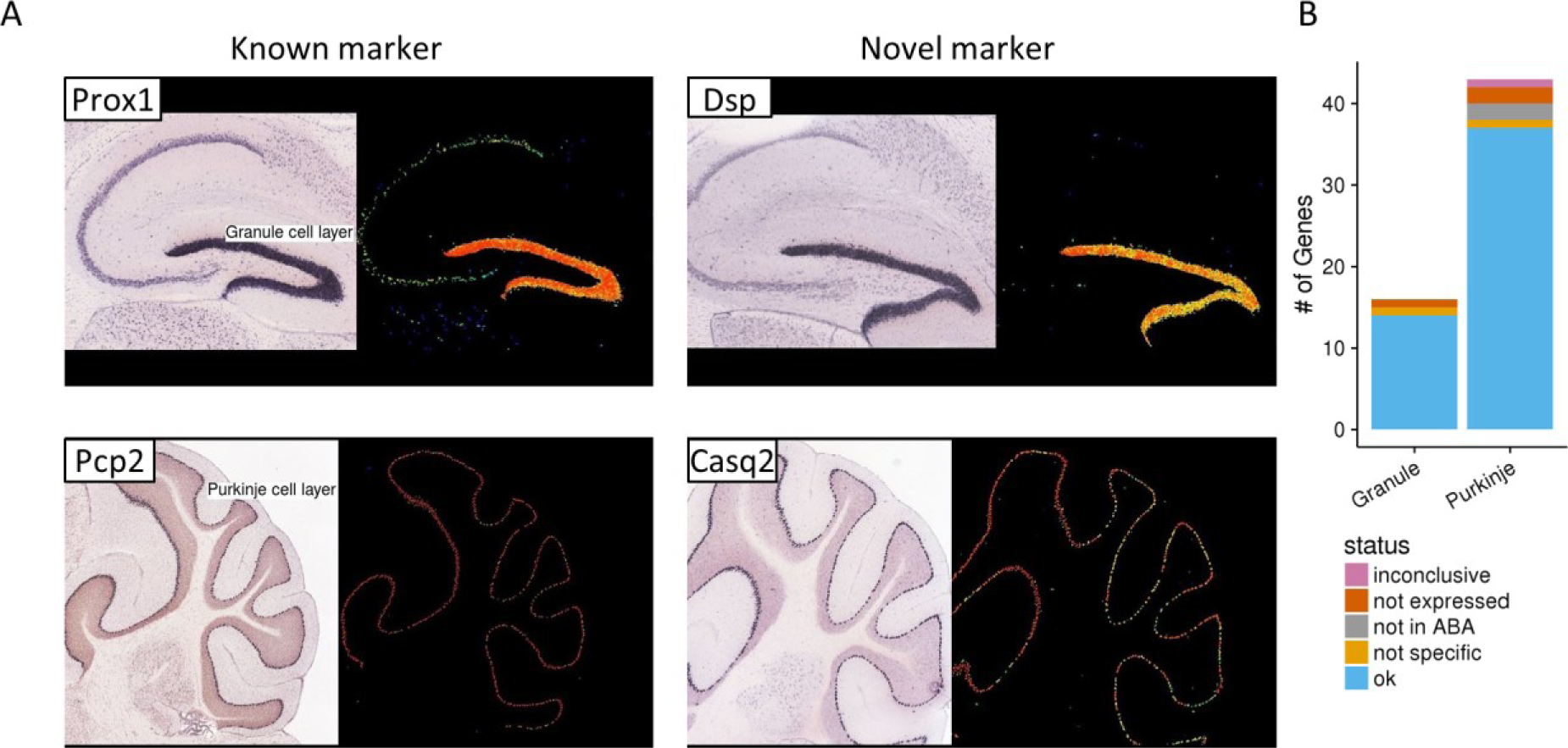
Validation of candidate markers using the Allen brain atlas. **(A)** In situ hybridization images from the Allen Brain Atlas. Rightmost panels show the location of the image in the brain according to the Allen Brain mouse reference atlas. Panels on the left show the ISH image and normalized expression level of known and novel dentate granule (upper panels) and Purkinje cell (lower panels) markers. **(B)** Validation status of marker genes detected for Purkinje and dentate granule cells. Figures used for validation and validation statuses of individual marker genes can be found in extended data (Figure 4-1,2,3,4).

The four markers for which no signal was detected (one marker of dentate gyrus granule cells and three markers of Purkinje cells) underwent additional scrutiny. For one of the markers of Purkinje cells (Eps8l2), the staining of cerebellar sections was inconsistent, with some sections showing no staining, some sections showing nonspecific staining and several sections showing the predicted localization. The three remaining genes had no signal in ABA ISH data brain-wide. We considered such absence or inconsistency of ISH signal inconclusive. Further analysis of these cases (one DG marker, three Purkinje) suggests that the ABA data is the outlier. As part of our marker selection procedure, Pter, the DG cell marker in question, was found to have high expression in granule cells both within NeuroExpresso and Hipposeq – a data set that is not used for primary selection of markers (see methods). In addition, Hipposeq indicates specificity to DG cells relative to the other neuron types in Hipposeq. For the Purkinje markers, specific expression for one gene (Sycp1) was supported by the work of Rong et al. (2004), who used degeneration of Purkinje cells to identify potential markers of these cells (20/43 Purkinje markers identified in our study were also among the list of potential markers reported by Rong et al.). We could not find data to further establish expression for the two remaining markers of Purkinje cells (Eps8l2 and Smpx). However, we stress that the transcriptomic data for Purkinje cells in our database are from five independent studies using different methodologies for cell purification, all of which support the specific expression of Eps8l2 and Smpx in Purkinje cells. Overall, through a combination of examination of ABA and other data sources, we were able to find confirmatory evidence of cell-type-specificity for 53/57 genes, with two false positives, and inconclusive findings for two genes.

We independently verified Cox6a2 as a marker of cortical fast spiking PV+ interneurons using triple label *in situ* hybridization of mouse cortical sections for Cox6a2, Pvalb and Slc32a1 (a pan-GABAergic neuronal marker) transcripts. As expected, we found that approximately 25% of all identified neurons were GABAergic (that is, Slc32a1 positive), while 46% of all GABAergic neurons were also Pvalb positive. 80% of all Cox6a2+ neurons were both Pvalb and Slc32a1 positive whereas Cox6a2 expression outside GABAergic cells was very low (1.65% of Cox6a2 positive cells), suggesting high specificity of Cox6a2 to PV+ GABAergic cells (Figure 5).

**Figure 5:**
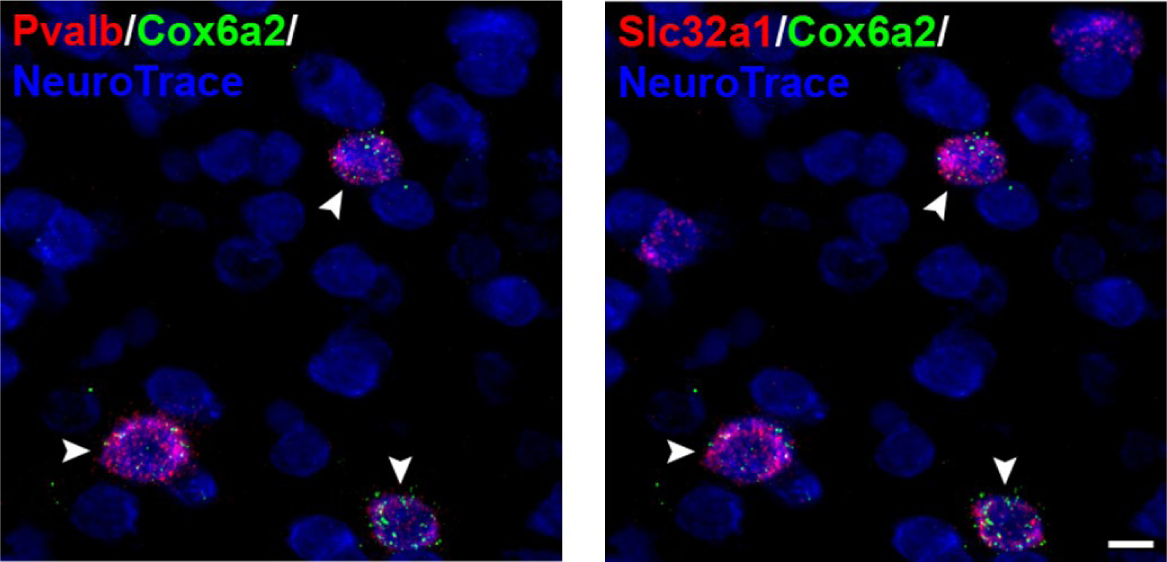
Single-plane image of mouse sensorimotor cortex labeled for Pvalb, Slc32a1, and Cox6a2 mRNAs and counterstained with NeuroTrace. Arrows indicate Cox6a2+ neurons. Bar = 10 µm.

### Verification of marker gene sets in single-cell RNA-seq data

As a further validation of our marker gene signatures, we analysed their properties in recently published single cell RNA-seq datasets derived from mouse cortex (Zeisel et al., 2015) and human cortex (Darmanis et al., 2015). We could not directly compare our MGSs to markers of cell type clusters identified in the studies producing these datasets since their correspondence to the cell types in NeuroExpresso was not clear. However, since both datasets represent a large number of individual cells, they are likely to include individual cells corresponding to the cortical cell types in our database. Thus, if our MGSs are cell type specific, and the corresponding cells are present in the single cell datasets, MGS should have a higher than random chance of being co-detected in the same cells, relative to non-marker genes. A weakness of this approach is that a failure to observe a correlation might be due to absence of the cell type in the data set rather than a true shortcoming of the markers. Overall, all MGSs for all cell types with the exception of oligodendrocyte precursor cells were successfully validated (p<0.001, Wilcoxon rank sum test) in both single cell datasets (<u>Table 3</u>).

**Table 3:** Coexpression of cortical MGSs in single cell RNA-seq data.

| Cell Types | Zeisel et al. (mouse) |  | Darmanis et al. (human) |  |
| --- | --- | --- | --- | --- |
|  | p-value | Gene Count | p-value | Gene Count |
| Endothelial | p<0.001 | 180 | p<0.001 | 157 |
| Astrocyte | p<0.001 | 282 | p<0.001 | 239 |
| Microglia | p<0.001 | 248 | p<0.001 | 201 |
| Oligodendrocyte | p<0.001 | 156 | p<0.001 | 201 |
| Oligodendrocyte precursors | 0.831 | 193 | 0.999 | 203 |
| FS Basket (G42) | p<0.001 | 26 | p<0.001 | 26 |
| Martinotti (GIN) | p<0.001 | 21 | p<0.001 | 20 |
| VIPReIn (G30) | p<0.001 | 43 | p<0.001 | 36 |
| Pyramidal | p<0.001 | 34 | p<0.001 | 27 |

Statistics are calculated by Wilcoxon rank sum test.

### NeuroExpresso as a tool for understanding the biological diversity and similarity of brain cells

One of the applications of NeuroExpresso is as an exploratory tool for exposing functional and biological properties of cell types. In this section, we highlight three examples we encountered: We observed high expression of genes involved in GABA synthesis and release (Gad1, Gad2 and Slc32a1) in forebrain cholinergic neurons, suggesting the capability of these cells to release GABA in addition to their cognate neurotransmitter acetylcholine (Figure 6A). Indeed, co-release of GABA and acetylcholine from forebrain cholinergic cells was recently demonstrated by Saunders et al. (2015). Similarly, the expression of the glutamate transporter Slc17a6, observed in thalamic (habenular) cholinergic cells suggests co-release of glutamate and acetylcholine from these cells, recently supported experimentally (Ren et al., 2011)) (Figure 6A). Surprisingly, we observed consistently high expression of Ddc (Dopa Decarboxylase), responsible for the second step in the monoamine synthesis pathway in oligodendrocyte cells (Figure 6B). This result is suggestive of a previously unknown ability of oligodendrocytes to produce monoamine neurotransmitters upon exposure to appropriate precursor, as previously reported for several populations of cells in the brain (Ren et al., 2016; Ugrumov, 2013). Alternatively, this finding might indicate a previously unknown function of Ddc. Lastly, we found overlap between the markers of spinal cord and brainstem cholinergic cells, and midbrain noradrenergic cells, suggesting previously unknown functional similarity between cholinergic and noradrenergic cell types. The common markers included Chodl, Calca, Cda and Hspb8, which were recently confirmed to be expressed in brainstem cholinergic cells (Enjin et al., 2010), and Phox2b, a known marker of noradrenergic cells (Pattyn et al., 1997).

**Figure 6:**
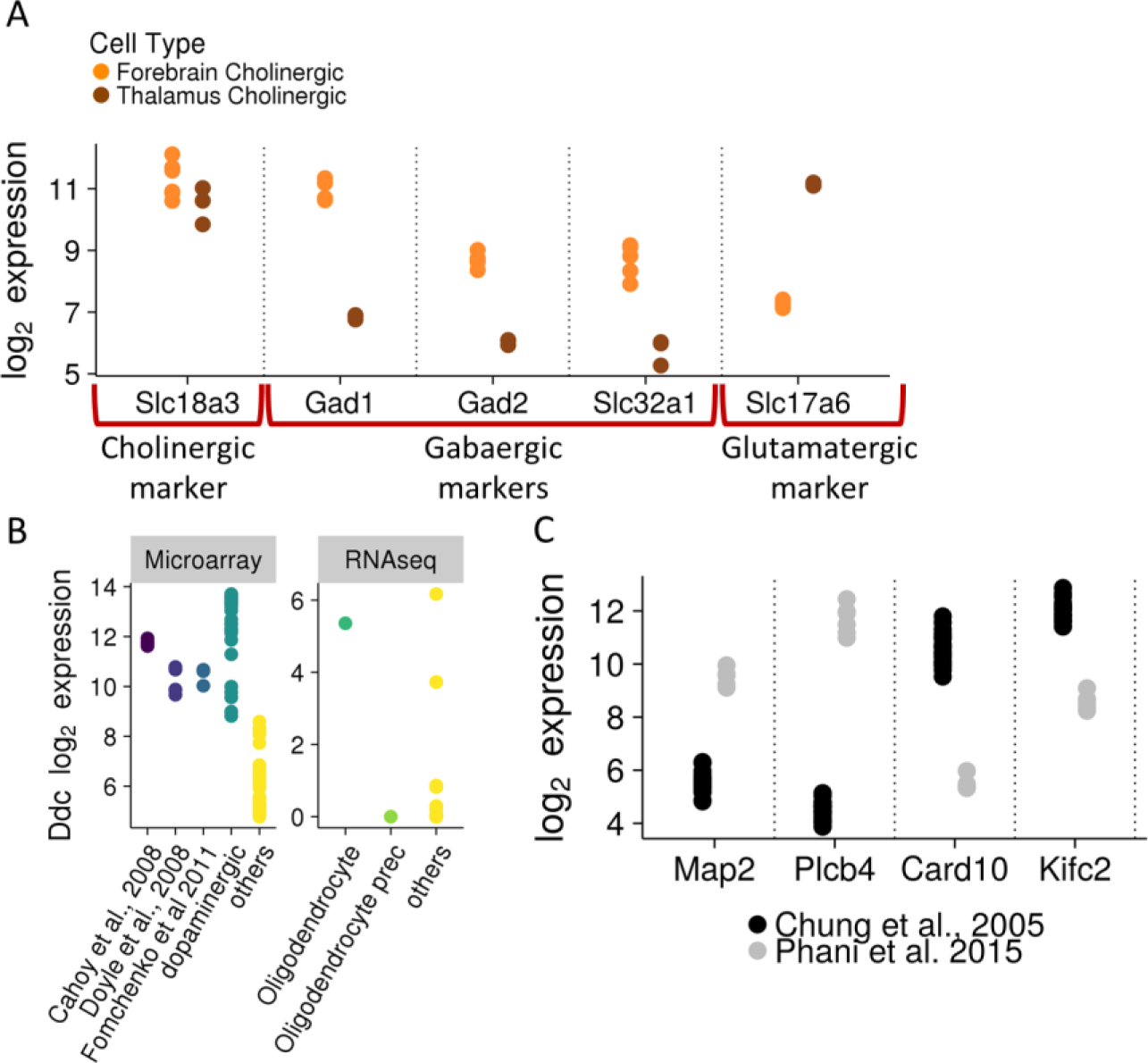
NeuroExpresso reveals gene expression patterns. **(A)** Expression of cholinergic, GABAergic and glutamatergic markers in cholinergic cells from forebrain and thalamus. Forebrain cholinergic neurons express GABAergic markers while thalamus (hubenular) cholinergic neurons express glutamatergic markers. **(B)** (Left) Expression of *Ddc* in oligodendrocyte samples from Cahoy et al., Doyle et al. and Fomchenko et al. datasets and in comparison to dopaminergic cells and other (non-oligodendrocyte) cell types from the frontal cortex in the microarray dataset. In all three datasets expression of *Ddc* in oligodendrocytes is comparable to expression in dopaminergic cells and is higher than in any of the other cortical cells. Oligodendrocyte samples show higher than background levels of expression across datasets. (Right) Ddc expression in oligodendrocytes, oligodendrocyte precursors, and other cell types from Tasic et al. single cell dataset. **(C)** Bimodal gene expression in two dopaminergic cell isolates by different labs. Genes shown are labeled as marker genes in the context of midbrain if the two cell isolates are labeled as different cell types.

### Marker Gene Profiles can be used to infer changes in cellular proportions in the brain

Marker genes are by definition cell type specific, and thus changes in their expression observed in bulk tissue data can represent either changes in the number of cells or cell type specific transcriptional changes (or a combination). Marker genes of four major classes of brain cell types (namely neurons, astrocytes, oligodendrocytes and microglia) were previously used to gain cell type specific information from brain bulk tissue data (Hagenauer et al., 2016; Kuhn et al., 2011; Ramaker et al., 2017; Sibille et al., 2008; Skene and Grant, 2016; P. P. C. Tan et al., 2013), and infer changes in cellular abundance. Following the practice of others, we applied similar approach to our marker genes, summarizing their expression profiles as the first principal component of their expression (see Methods) (Chikina et al., 2015; Westra et al., 2015; Xu et al., 2013). We refer to these summaries as Marker Gene Profiles (MGPs).

In order to validate the use of MGPs as surrogates for relative cell type proportions, we used bulk tissue expression data from conditions with known changes in cellular proportions. Firstly, we calculated MGPs for human white matter and frontal cortex using data collected by (Trabzuni et al., 2013). Comparing the MGPs in white vs. grey matter, we observed the expected increase in oligodendrocyte MGP, as well as increase in oligodendrocyte progenitor cell, endothelial cell, astrocyte and microglia MGPs, corroborating previously reported higher number of these cell types in white vs. grey matter (Gudi et al., 2009; Ogura et al., 1994; Williams et al., 2013). We also observed decrease in MGPs of all neurons, corroborating the low neuronal cell body density in white vs. grey matter (Figure 7A, <u>Table 4</u>).

**Table 4:**
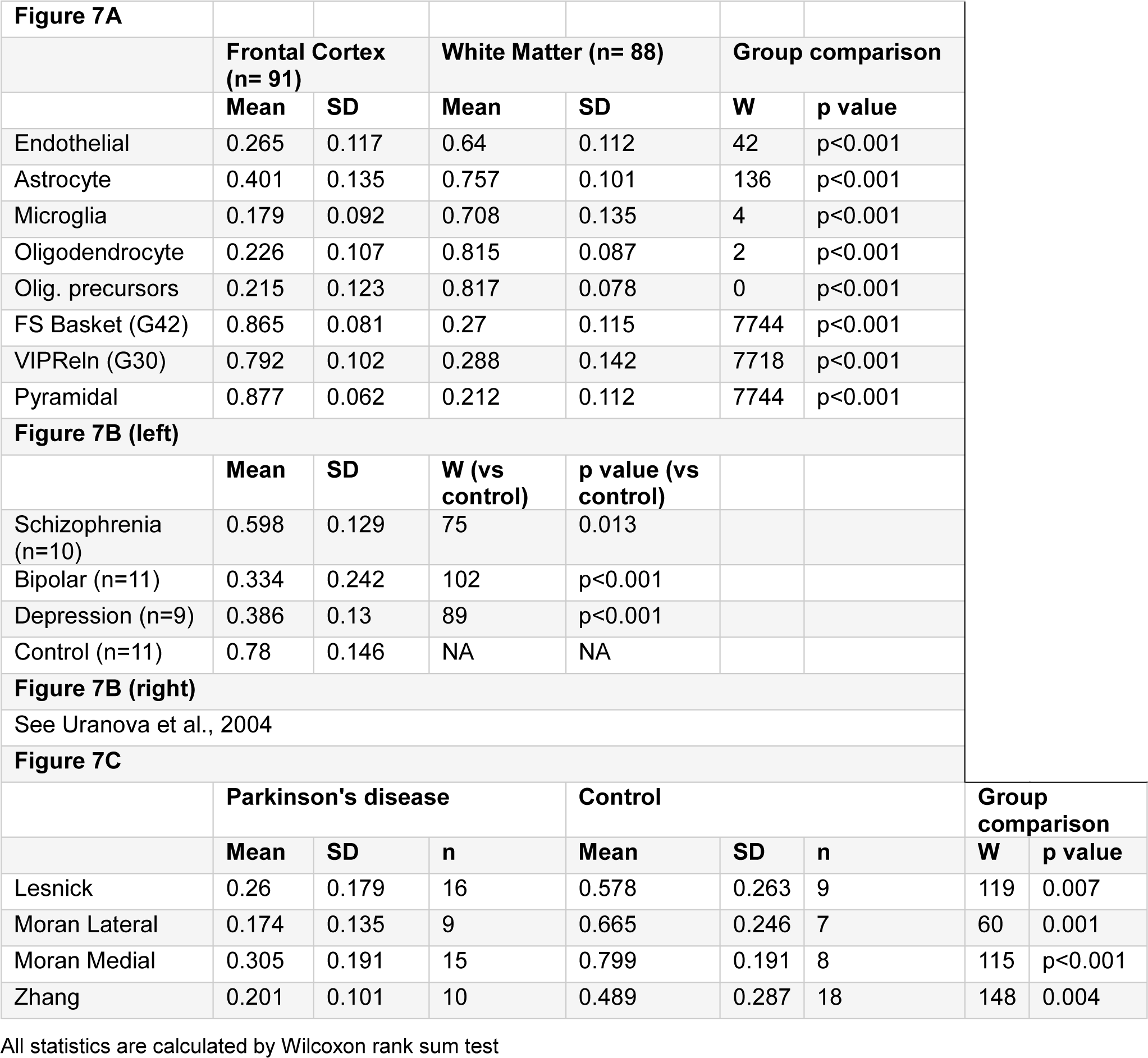
Summaries of statistical analyses.

| Figure 7A |  |  |  |  |  |  |  |  |
| --- | --- | --- | --- | --- | --- | --- | --- | --- |
|  | Frontal Cortex (n= 91) |  | White Matter (n= 88) |  | Group comparison |  |  |  |
|  | Mean | SD | Mean | SD | W | p value |  |  |
| Endothelial | 0.265 | 0.117 | 0.64 | 0.112 | 42 | p<0.001 |  |  |
| Astrocyte | 0.401 | 0.135 | 0.757 | 0.101 | 136 | p<0.001 |  |  |
| Microglia | 0.179 | 0.092 | 0.708 | 0.135 | 4 | p<0.001 |  |  |
| Oligodendrocyte | 0.226 | 0.107 | 0.815 | 0.087 | 2 | p<0.001 |  |  |
| Olig. precursors | 0.215 | 0.123 | 0.817 | 0.078 | 0 | p<0.001 |  |  |
| FS Basket (G42) | 0.865 | 0.081 | 0.27 | 0.115 | 7744 | p<0.001 |  |  |
| VIPReIn (G30) | 0.792 | 0.102 | 0.288 | 0.142 | 7718 | p<0.001 |  |  |
| Pyramidal | 0.877 | 0.062 | 0.212 | 0.112 | 7744 | p<0.001 |  |  |
| Figure 7B (left) |  |  |  |  |  |  |  |  |
|  | Mean | SD | W (vs control) | p value (vs control) |  |  |  |  |
| Schizophrenia (n=10) | 0.598 | 0.129 | 75 | 0.013 |  |  |  |  |
| Bipolar (n=11) | 0.334 | 0.242 | 102 | p<0.001 |  |  |  |  |
| Depression (n=9) | 0.386 | 0.13 | 89 | p<0.001 |  |  |  |  |
| Control (n=11) | 0.78 | 0.146 | NA | NA |  |  |  |  |
| Figure 7B (right) |  |  |  |  |  |  |  |  |
| See Uranova et al., 2004 |  |  |  |  |  |  |  |  |
| Figure 7C |  |  |  |  |  |  |  |  |
|  | Parkinson's disease |  |  | Control |  |  | Group comparison |  |
|  | Mean | SD | n | Mean | SD | n | W | p value |
| Lesnick | 0.26 | 0.179 | 16 | 0.578 | 0.263 | 9 | 119 | 0.007 |
| Moran Lateral | 0.174 | 0.135 | 9 | 0.665 | 0.246 | 7 | 60 | 0.001 |
| Moran Medial | 0.305 | 0.191 | 15 | 0.799 | 0.191 | 8 | 115 | p<0.001 |
| Zhang | 0.201 | 0.101 | 10 | 0.489 | 0.287 | 18 | 148 | 0.004 |

All statistics are calculated by Wilcoxon rank sum test

A more specific form of validation was obtained from a pair of studies done on the same cohort of subjects, with one study providing expression profiles (study 2 from SMRI microarray database, see Methods) and another providing stereological counts of oligodendrocytes (Uranova et al., 2004), for similar brain regions. We calculated oligodendrocyte MGPs based on the expression data and compared the results to experimental cell counts from Uranova et al. (2004). The MGPs were consistent with the reduction of oligodendrocytes observed by Uranova et al. in schizophrenia, bipolar disorder and depression patients. (Figure 7B, <u>Table 4;</u> direct comparison between MGP and experimental cell count at a subject level was not possible, as Uranova et al. did not provide subject identifiers corresponding to each of the cell count values). To further assess and demonstrate the ability of MGPs to correctly represent cell type specific changes in neurological conditions, we calculated dopaminergic profiles of substantia nigra samples in three expression data sets of Parkinson’s disease (PD) patients and controls from Moran et al. (2006) (GSE8397), Lesnick et al. (2007) (GSE7621) and Zhang et al. (2005) (GSE20295). We tested whether the well-known loss of dopaminergic cells in PD could be detected using our MGP approach. MGP analysis correctly identified reduction in dopaminergic cells in substantia nigra of Parkinson’s disease patients (Figure 7C, <u>Table 4</u>).

## Discussion

### Cell type specific expression database as a resource for neuroscience

We present NeuroExpresso, a rigorously curated database of brain cell type specific gene expression data (<u>www.neuroexpresso.org)</u>, and demonstrate its utility in identifying cell-type markers and in the interpretation of bulk tissue expression profiles. To our knowledge, NeuroExpresso is the most comprehensive database of expression data for identified brain cell types. The database will be expanded as more data become available.

NeuroExpresso allows simultaneous examination of gene expression associated with numerous cell types across different brain regions. This approach promotes discovery of cellular properties that might have otherwise been unnoticed or overlooked when using gene-by-gene approaches or pathway enrichment analysis. For example, a simple examination of expression of genes involved in biosynthesis and secretion of GABA and glutamate, suggested the co-release of these neurotransmitters from forebrain and habenular cholinergic cells, respectively.

Studies that aim to identify novel properties of cell types can benefit from our database as an inexpensive and convenient way to seek novel patterns of gene expression. For instance, our database shows significant bimodality of gene expression in dopaminergic cell types from the midbrain (Figure 6C). The observed bimodality might indicate heterogeneity in the dopaminergic cell population, which could prove a fruitful avenue for future investigation. Another interesting finding from NeuroExpresso is the previously unknown overlap of several markers of motor cholinergic and noradrenergic cells. While the overlapping markers were previously shown to be expressed in spinal cholinergic cells, to our knowledge their expression in noradrenergic (as well as brain stem cholinergic) cells was previously unknown.

NeuroExpresso can be also used to facilitate interpretation of genomics and transcriptomics studies. Recently (Pantazatos et al., 2016) used an early release of the databases to interpret expression patterns in the cortex of suicide victims, suggesting involvement of microglia. Moreover, this database has further applications beyond the use of marker genes, such as understanding the molecular basis of cellular diversity (Tripathy et al., 2017).

Importantly, NeuroExpresso is a cross-laboratory database. A consistent result observed across several studies raises the certainty that it represents a true biological finding rather than merely an artefact or contamination with other cell types. This is specifically important for unexpected findings such as the expression of Ddc in oligodendrocytes (Figure 6B).

### Validation of cell type markers

To assess the quality of the marker genes, a subset of our cell type markers was validated by in situ hybridization (Cox6a2 as a marker of fast spiking basket cells, and multiple Purkinje and DG cell markers). Further validation was performed with computational methods in independent single cell datasets from mouse and human. This analysis validated all cortical gene sets except Oligodendrocyte precursors (OP). In their paper, Zeisel et al. (2015) stated that none of the oligodendrocyte sub-clusters they identified were associated with oligodendrocyte precursor cells, which likely explains why we were not able to validate the OP MGP in their dataset. The Darmanis dataset however, is reported to include oligodendrocyte precursors (18/466 cells) (Darmanis et al., 2015), but again our OPC MGP did not show good validation. In this case the reason for negative results could be changes in the expression of the mouse marker gene orthologs in human, possibly reflecting functional differences between the human and mouse cell types (Shay et al., 2013; Zhang et al., 2016). Further work will be needed to identify a robust human OPC signature. However, since most MGSs did validate between mouse and human data, it suggests that most marker genes preserve their specificity despite cross-species gene expression differences.

### Improving interpretation of bulk tissue expression profiles

Marker genes can assist with the interpretation of bulk tissue data in the form of marker gene profiles (MGPs). A parsimonious interpretation of a change in an MGP is a change in the relative abundance of the corresponding cell type. Similar summarizations of cell type specific genes were previously used to analyse gene expression (Chikina et al., 2015; Newman et al., 2015; Westra et al., 2015; Xu et al., 2013) and methylation data (Jones et al., 2017; Shannon et al., 2017). Since our approach focuses on the overall trend of a MGS expression level, it should be relatively insensitive to expression changes in a subset of these genes. Still, we prefer to refer the term “marker gene profile” rather than “cell type proportions”, to emphasize the indirect nature of the approach.

Our results show that MGPs based on NeuroExpresso marker gene sets (MGSs) can reliably recapitulate relative changes in cell type abundance across different conditions. Direct validation of cell count estimation based on MGSs in human brain was not feasible due to the unavailability of cell counts coupled with expression data. Instead, we compared oligodendrocyte MGPs based on a gene expression dataset available through the SMRI database to experimental cell counts taken from a separate study (Uranova et al., 2004) of the same cohort of subjects and were able to recapitulate the reported reduction of oligodendrocyte proportions in patients with schizophrenia, bipolar disorder and depression. Based on analysis of dopaminergic MGPs we were also able to capture the well-known reduction in dopaminergic cell types in PD patients.

### Limitations and caveats

While we took great care in the assembly of NeuroExpresso, there remain a number of limitations and room for improvement. First, the NeuroExpresso database was assembled from multiple datasets, based on different mouse strains and cell type extraction methodologies, which may lead to undesirable heterogeneity. We attempted to reduce inter-study variability by combined pre-processing of the raw data and normalization. However, due to insufficient overlap between cell types represented by different studies, many of the potential confounding factors such as age, sex and methodology could not be explicitly corrected for. Thus, it is likely that some of the expression values in NeuroExpresso may be affected by confounding factors. While our confidence in the data is increased when expression signals are robust across multiple studies, many of the cell types in NeuroExpresso are represented by a single study. Hence, we advise that small differences in expression between cell types as well as previously unknown expression patterns based on a single data source should be treated with caution. In our analyses, we address these issues by enforcing a stringent set of criteria for the marker selection process, reducing the impact of outlier samples, ignoring small changes in gene expression and validating the results in external data. However, it must be noted that it was not possible validate our markers for all cell types and brain regions.

An additional limitation of our study is that the representation for many of the brain cell types is still lacking in the NeuroExpresso database. Therefore, despite our considerable efforts to ensure cell type-specificity of the marker genes, we cannot rule out the possibility that some of them are also expressed in one or more of the non-represented cell types. This problem is partially alleviated in cortex due to the inclusion of single cell data. As more such datasets become available, it will be easier to create a more comprehensive database. A related problem to the coverage of cell types in NeuroExpresso lies in the definition of the term “cell type”. Most cell types represented in NeuroExpresso are heterogeneous populations. For instance, fast-spiking basket cells as defined by microarray data matches 5 distinct clusters identified by Tasic et al. (2016) based on single cell RNA sequencing data. By considering them as a single cell type, we lose the ability to detect unique properties of the individual clusters. Heterogeneity also may reduce the confidence we have in our marker genes. If a selected marker is expressed in a subtype of another cell type, this will not be noticed in pooled expression data as the signal will be suppressed by other subtypes that do not express the gene. We hope to remedy this problem with increased availability of single cell data in the future. Where inter-cell type variability ends and new cell type begins is an ongoing discussion in the field. For the purposes of this study, we tried to ensure that cell types we define are accepted and studied by a portion of the community, and that the expression profiles of the cell types were distinct enough to allow marker gene identification. The data we make available to other researchers may be portioned into finer cell types or grouped together into more broad cell type groups depending on the aims of the researchers.

Finally, it must be noted that while we aim to infer changes in cell type abundance with MGPs, we do not attempt to estimate the cell type proportions themselves even though many established deconvolution methods do accomplish this using databases of expression profiles (Chikina et al., 2015; Grange et al., 2014; Newman et al., 2015). These approaches operate on the assumption that the absolute expression levels of genes will be conserved across the cell types in the reference database and cell types that make up the whole tissue sample. In our work, we avoid these approaches because our database (mouse cell types) and the whole tissue samples we analyse (human brain tissue) come from different species which may cause changes in gene expression, while marker genes are more likely to be conserved.

In summary, we believe that NeuroExpresso is a valuable resource for neuroscientists. We identified numerous novel markers for 36 major cell types and used them to estimate cell type profiles in bulk tissue data, demonstrating high correlation between our estimates and experiment-based cell counts. This approach can be used to reveal cell type specific changes in whole tissue samples and to re-evaluate previous analyses on brain whole tissues that might be biased by cell type-specific changes. Information about cell type-specific changes is likely to be very valuable since conditions like neuron death, inflammation, and astrogliosis are common hallmarks of in neurological diseases.

## Acknowledgements

We thank Ken Sugino for providing access to raw CEL files for Purkinje and TH+ cells from locus coeruleus, Chee Yeun Chung for providing access to raw CEL files for dopaminergic cells, Dean Attali for providing insight on the usage of the Shiny platform, the Pavlidis lab for their inputs to the project during its development and Rosemary McCloskey for aid in editing the manuscript.

## Extended Data

### Figure 4B

Figure 4-1: Expression of dentate granule cell markers discovered in the study in Allen Brain Atlas mouse brain in situ hybridization database. The first gene is Prox1, a known marker of dentate granule cells. The intensity is color-coded to range from blue (low expression intensity), through green (medium intensity) to red (high intensity). All images except Ogn is taken from the sagittal view. Ogn is taken from the coronal view.

Figure 4-2: Expression of Purkinje markers discovered in the study in Allen Brain Atlas mouse brain in situ hybridization database. The first gene is Pcp2, a known marker of Purkinje cells. The intensity is color-coded to range from blue (low expression intensity), through green (medium intensity) to red (high intensity).} All images are taken from the sagittal view.

Figure 4-3: Validation status of dentate granule cell markers.

Figure 4-4: Validation status of Purkinje cell markers.

### neuroExpressoAnalysis-master.zip

Code for data acquisition, analysis and generation of all figures.

### markerGeneProfile-master.zip

R package to perform marker gene profile estimations on whole tissue expression data and to select marker genes from cell type specific expression data.

### homologene-master.zip

R package to find gene homologues across species.

